# The developmental basis for scaling of mammalian tooth size

**DOI:** 10.1101/2022.12.03.518730

**Authors:** Mona M. Christensen, Outi Hallikas, Rishi Das Roy, Vilma Väänänen, Otto E. Stenberg, Teemu J. Häkkinen, Jean-Christophe François, Robert J. Asher, Ophir D. Klein, Martin Holzenberger, Jukka Jernvall

## Abstract

When evolution leads to differences in body size, organs generally scale along. A well-known example of the tight relationship between organ and body size is the scaling of mammalian molar teeth. To investigate how teeth scale during development and evolution, we compared molar development in mouse and rat from initiation through final size. Whereas the linear dimensions of the rat first lower molar are twice that of the mouse molar, their shapes are largely the same. We found that scaling of the molars starts early, and that the rat molar is patterned equally as fast but in a larger size than the mouse molar. Using transcriptomics, we discovered that a known regulator of body size, insulin-like growth factor 1 (*Igf1*), is more highly expressed in the rat molars compared to the mouse molars. *Ex vivo* and *in vivo* mouse models demonstrated that modulation of the IGF pathway reproduces several aspects of the observed scaling process. Furthermore, analysis of IGF1-treated mouse molars and computational modeling indicate that IGF signaling scales teeth by simultaneously enhancing growth and by inhibiting the cusp patterning program, thereby providing a relatively simple mechanism for scaling teeth during development and evolution. Finally, comparative data from shrews to elephants suggest that this scaling mechanism regulates the minimum tooth size possible, as well as the patterning potential of large teeth.

## INTRODUCTION

Body size evolution causes fundamental changes in an organism’s ecology and physiology (Peters, 1983). Changes in body size have been well documented for multiple taxonomic groups (Smith et al., 2016), and these changes in overall size are typically tightly linked to the scaling of individual body parts and organs (Damuth & MacFadden, 1990). The mammalian molar tooth is an example of an organ that scales with body size. This scaling link is so strong that, within evolutionary lineages, highly accurate estimates of body size can be made from simple linear measures of molar teeth (Damuth & MacFadden, 1990; Hopkins, 2018). The use of linear measurements to estimate body size is made possible by the relatively shape-invariant scaling of molars within mammalian lineages (Damuth & MacFadden, 1990; Copes & Schwartz, 2010; Hopkins, 2018). As a result, the fossil record of molars forms much of the basis for the reconstructions of the body size dynamics in mammalian evolution (Gingerich, 1982; Alroy, 1998; Smith et al., 2010; D’Ambrosia et al., 2017).

Despite an increasing understanding of the molecular mechanisms of shape and overall size regulation (Parker, 2011; Boulan & Leopold, 2021; Harmansa & Lecuit, 2021; Wu & Guan, 2020), it remains unknown how evolutionary changes in organ size are achieved while keeping shape and proportions constant. Given the central role of mammalian molars in estimating body size, we set out to investigate how the shape-invariant scaling is realized during tooth development. We took advantage of the fact that two mammalian species used in developmental research, the mouse (*Mus musculus*) and the rat (*Rattus norvegicus*), provide an example of divergent body and molar tooth size but relatively similar molar tooth shape (**Fig. 1A**). This shape-invariant scaling allows us to focus on size alone, without the pervasive effects of shape differences during development (Jernvall et al., 2000).

**Figure 1.**
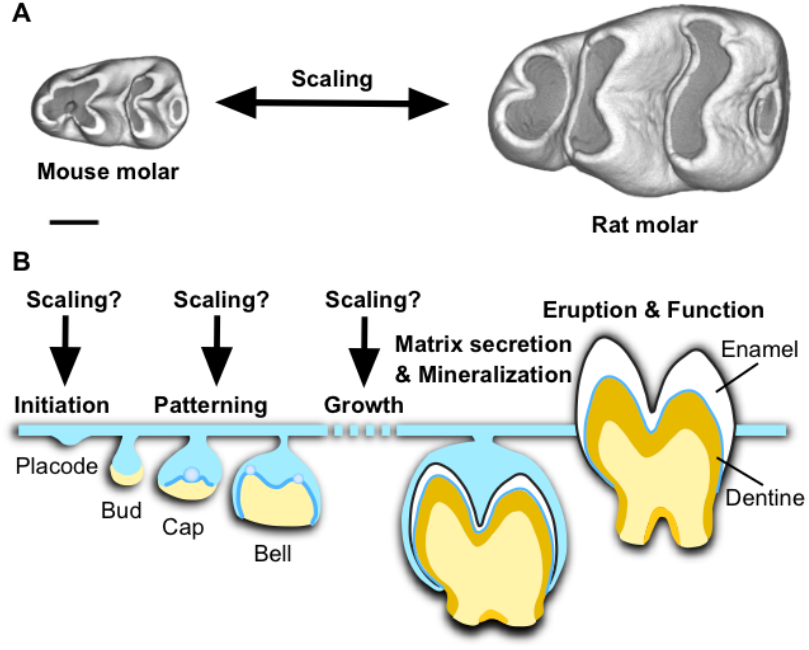
Determining when teeth are scaled during development. **A**, First lower molars of the mouse and the rat are similar in overall shape, but the rat molar is two times larger in linear dimensions. Occlusal views, anterior to the left, buccal to the top. Scale bar, 500 µm. **B**, When and how during tooth development the scaling process occurs is not known. Tooth development is regulated by the interactions between the epithelial (blue) and mesenchymal (yellow) tissues. After mineralization and eruption, crown shape cannot be remodeled.

## RESULTS

### Molar scaling begins during the placode stage

As a first step, we established when the size differences between mouse and rat molars begin to appear during development. Specifically, we asked whether size differences become visible already during the patterning of cusps, or whether rat molars achieve their larger size through growth after patterning (**Fig. 1B**). Whereas the patterning process of mammalian teeth is well known to integrate inductive signaling and growth (Harjunmaa et al., 2014), it is not known whether and how scaling might be involved.

To pinpoint the onset of scaling, we compared molar development of the mouse and the rat chronologically by starting from the dental placodes. These are the earliest individualized dental structures to form when the epithelium begins to invaginate into the underlying mesenchyme. Because it is difficult to reliably delineate the size of the epithelial placode morphologically, we used *in situ* hybridization to detect gene expression of two epithelial markers, forkhead box I3 (*Foxi3*) and sonic hedgehog (*Shh*). *Foxi3* expression encompasses the entire placodal epithelium (Shirokova et al., 2013), and *Shh* is expressed within the placode in the early signaling centers, the initiation knots (Mogollon et al., 2021).

When multiple placodes are examined, *Foxi3* expression domains overlap in size between the species (**Fig. 2A**, *p* = 0.1626, all *p*-values determined using two-tailed randomization test, **Table S1)**. This suggests that the overall sizes of the placodal epithelia are similar in the species. However, *Shh* expression domain, which is upregulated within the placode in the initiation knot (Mogollon et al., 2021), is slightly larger in the rat than in the mouse (**Fig. 2B**, *p* = 0.0008, **Table S1**). Formation of the initiation knot marks the beginning of the transition from placode to bud stage, and this step appears to also mark the beginning of scaling.

**Figure 2.**
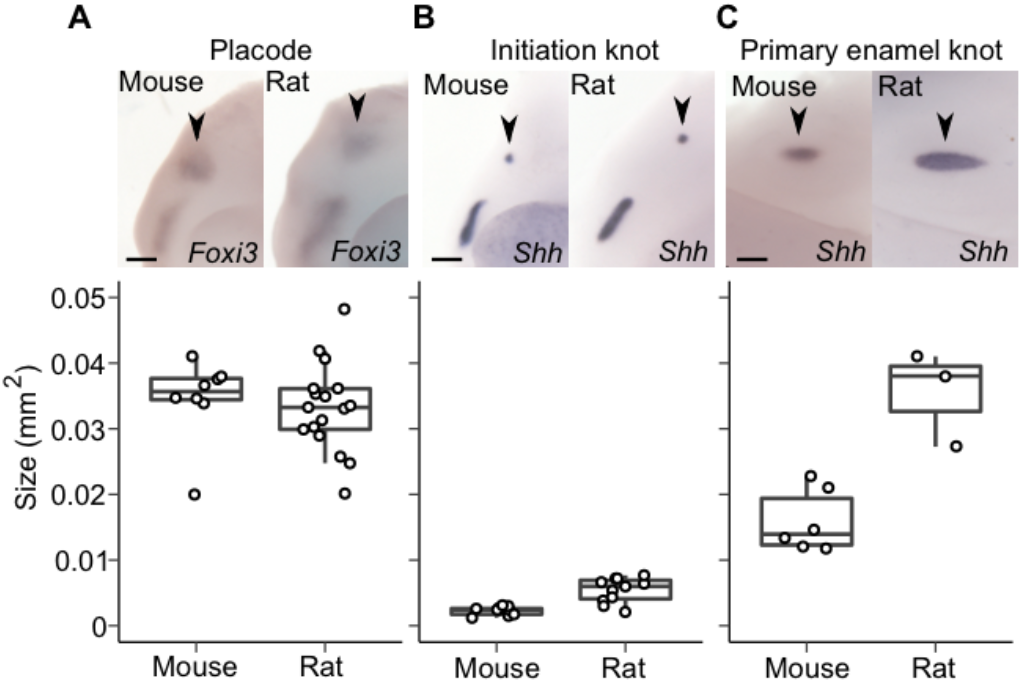
Tooth scaling begins during the placode stage of molar development. **A**, The epithelial placodes (black arrowheads) are similar in size in the rat (*n* = 17) and mouse molar, visualized using *Foxi3* expression (*n* = 8, randomization test *p* = 0.1626). **B**, The initiation knots (black arrowheads, visualized using *Shh* expression) are larger in the rat (*n* = 11) than in the mouse (*n* = 9, *p* = 0.0008). **C**, The primary enamel knots (black arrowheads, visualized using *Shh* expression) are larger in the rat (*n* = 3) than in the mouse (*n* = 6, *p* = 0.0131. Boxes enclose 50% of observations, the horizontal bar denotes the median, and whiskers extend to last values within 1.5 interquartiles. For the images, anterior is to the left, buccal to the top. Scale bars, 200 µm.

Examining the expression patterns in more detail shows that the size difference between the mouse and rat is driven by an increasing difference along the longitudinal axis (**Table S1**). This difference becomes more pronounced when the primary enamel knot appears two days after the placode stage in both species (day E14 and E16 in the mouse and the rat, respectively). The primary enamel knot is an epithelial signaling center that forms towards the end of the bud stage when the invaginated epithelial bud starts to grow lateral folds called the cervical loops. Cervical loop growth marks the onset of the cap stage, during which tooth crown morphogenesis begins. The rat primary enamel knot, detected with *Shh* expression, is roughly twice as large in area as that of the mouse (**Fig. 2C, Table S1**), suggesting a marked difference in signaling activity between the species at the onset of crown formation. Overall, scaling of tooth size appears to start before the active patterning of cusps.

### Molar scaling encompasses all the patterning stages

To examine whether scaling is a significant factor affecting growth during patterning, we analyzed developing crown morphologies. From the cap stage onwards, size measurements can be carried out using three-dimensional reconstructions. We used soft-tissue µCT imaging to reconstruct both the size and shape of the growing molars (Methods). To quantify the overall progression of patterning in which individual cusps become gradually identifiable, we used Orientation Patch Count (OPC) to measure surface complexity (Evans et al., 2007).

Aligning the growth series of mouse and rat molars based on days after placode initiation makes it apparent that rat molars grow substantially faster (**Fig. 3A**). When plotted (using mm^2^, **Fig. 3B, Table S2**), both molars appear to achieve their final sizes within about 10 days after the placode stage. Logistic growth models fitted for the data suggest that the inflection points, after which the growth begins to slow down, occur close to seven days after the placode stage (day E19 and E21 for the mouse and rat, respectively, **Fig. 3B**). Considering that the stage with final number of main cusps is separated by seven days from the placode stage in both species (**Fig. 3A)**, scaling appears to encompass all the stages of cusp patterning. Indeed, in contrast to the pronounced differences in size, OPC values show largely similar increases of topographic complexity in the two species (**Fig. 3C, D, Table S2**). The slightly higher OPC values of the rat molar reflect its more distinct anterior part of the crown (anteroconid) and an additional distobuccal cusp (arrowheads in **Fig. 3A)**. The inflection points of increase in complexity precede those of the increase in size by 2.5 and 1.4 days for the mouse and rat, respectively (**Fig. 3D**), further indicating that patterning is embedded within the scaling process of teeth.

**Figure 3.**
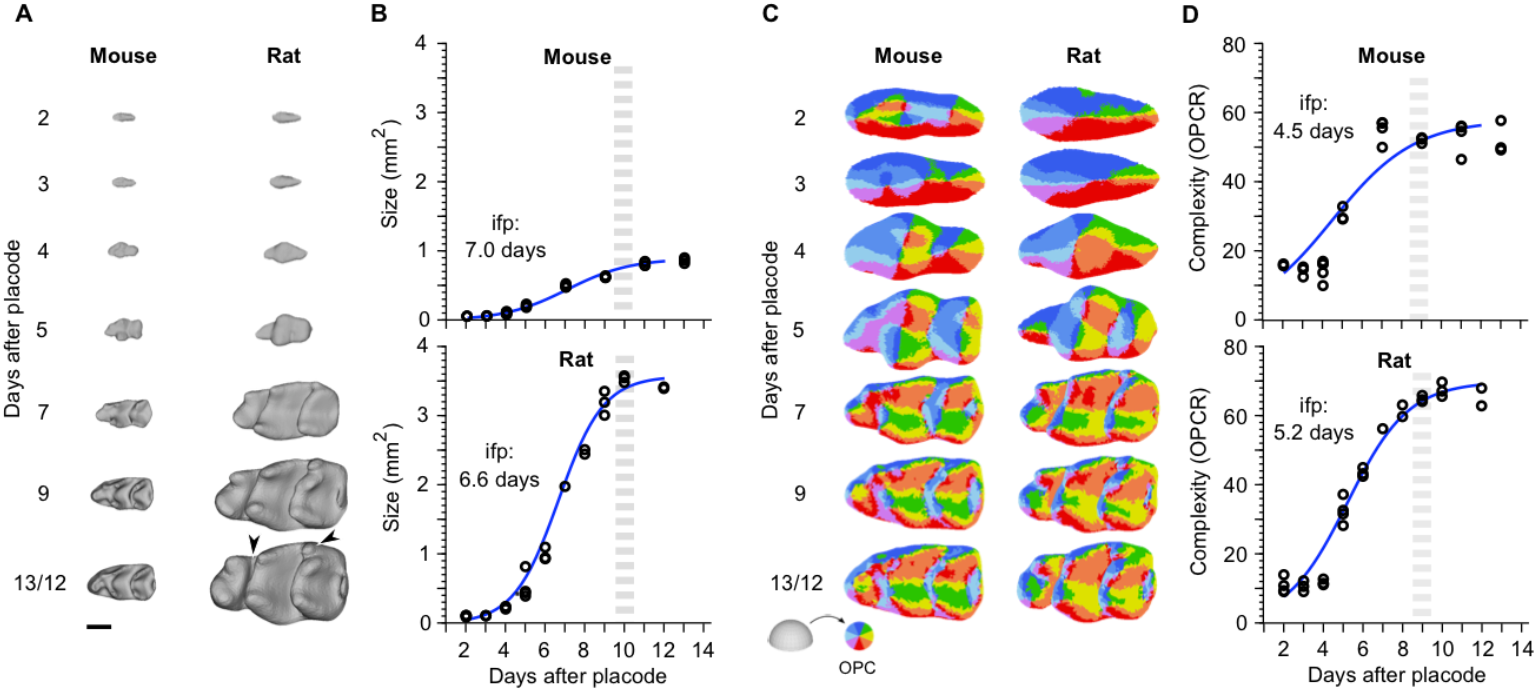
Despite accelerated growth in size, rat molar patterning is similar to mouse molar patterning. **A**, 3D-reconstructions show rat molars becoming progressively larger throughout development (*n* = 26 and 28 for mouse and rat). **B**, Despite the much faster growth of the rat molar, logistic curves fitted to the areas indicate comparable time points for the onset of growth deceleration (inflection point, ifp) and reaching of the final tooth size (grey dashed line). For logistic curve to calculate the size *S(t) = K/*1+*Ae*^*-kt*^; *K, A*, and *k* are 0.9, 80.9, and 0.62 for the mouse, and 33.6, 350, and 0.88 for the rat, respectively. **C**, OPC maps of dental complexity show generally comparable progression of patterning. **D**, Compared to the size, logistic curves fitted to the OPCR values are relatively similar between the species, the inflection point being slightly earlier in the mouse. For logistic curve to calculate the OPC *O(t) = K/*1+*Ae*^*-kt*^; *K, A*, and *k* are 57.8, 8.6, and 0.48 for the mouse, and 69.8, 29.6, and 0.65 for the rat, respectively. The higher OPCR values of the rat molar reflect its more distinct anteroconid and an additional distobuccal cusp (arrowheads in **A**). Anterior to the left, buccal to the top. Scale bar, 500 µm in (**A**).

Taken together, these results point to largely comparable rates of shape development between the mouse and the rat molars, although the teeth themselves increase in size at very different rates. A major implication of this observation is that the patterning happens in tissue domains that differ in size. This in turn indicates that the patterning process itself scales.

### Scaling of patterning involves changes in spacing of signaling centers

Morphological appearance of cusps is preceded up to one day by the formation of transient signaling centers, called the secondary enamel knots, that differentiate at the locations of the future cusps (Jernvall et al., 2000). Because the OPC analysis shows that patterning occurs in larger size in the rat (**Fig. 3**), we used the expression of fibroblast growth factor 4 (*Fgf4*) to examine whether the spacing of the secondary enamel knots also differs between the mouse and rat (**Fig. 4A**). The results confirm that during the patterning stage the rat molar is not only larger than the mouse molar, but also that the secondary enamel knots are more spread apart (**Fig. 4A, b**, *p* = 0.0001 for both size and patterning, **Table S3**). Thus, the scaling of tooth size appears to be linked to the dynamics of patterning regulation, and not, for example, to cell size differences between the species (**Fig. S1**).

**Figure 4.**
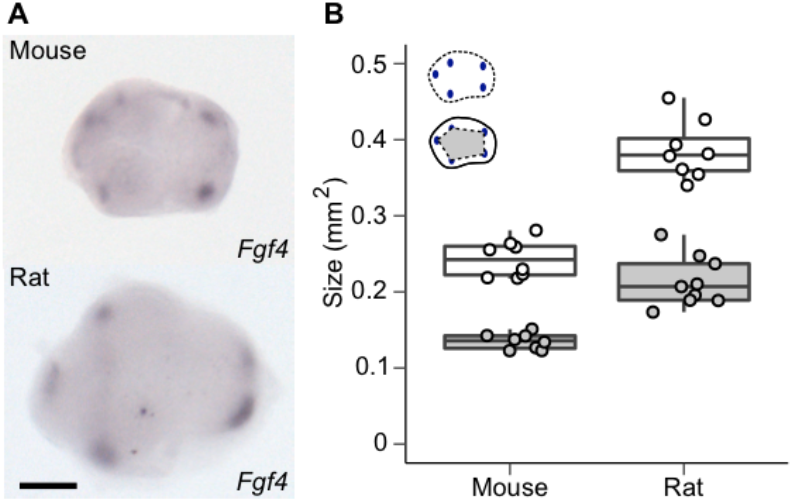
Scaling of patterning involves changes in tooth size and spacing of signaling centers. **A**, Secondary enamel knots visualized using *in situ* hybridization of *Fgf4* expression in the mouse and rat molar. **B**, The rat molar is larger (size shown with white points, *n* = 8), and the secondary enamel knots are more spread apart (patterning area shown with grey points, *n* = 9) than in the mouse molar (*n* = 8 for both measurements). All *p*-values are 0.0001. Boxes enclose 50% of observations, the horizontal bar denotes the median, and whiskers extend to last values within 1.5 interquartiles. Anterior to the left, buccal to the top. Scale bar, 200 µm.

### Modifying IGF1 signaling is sufficient to scale both size and patterning

Next, we examined how signaling and regulation of proliferation are integrated to scale teeth. To identify molecular mechanisms that could explain the scaling of both tooth size and patterning, we first used RNA sequencing (RNAseq) to compare gene expression between the two species. We performed RNAseq analyses for mouse and rat molars that were one, three, and four days from the placode stage (corresponding to bud, late cap, and bell stages, Methods). Although the overall expression levels of genes required for normal tooth development are highly comparable between the species (Hallikas et al., 2021), we found that many genes of the insulin-like growth factor (*Igf*) pathway were expressed at higher levels in the rat than in the mouse molar (**Fig. 5A, Table S4**). In particular, *Igf1* was consistently expressed at much higher levels in the rat (**Fig. 5A, Table S4**), and *Igf2*, which also functions through the IGF1 receptor, showed higher expression levels in the later stages of the rat molar development (**Table S4**). Moreover, several of the genes encoding IGF-binding proteins (IGFBPs) that modulate local IGF1-signaling were highly expressed in the rat molar (**Table S4**). The IGF1 receptor mediated pathway is required for various aspects of tissue growth, such as proliferation and survival (Dupont & Holzenberger, 2003; LeRoith et al., 2021), and it is well established as a regulator of body size in dogs, humans and mice (Woods et al., 1996; Sjögren et al., 1999; Sutter et al., 2007). Whereas IGF1 functions postnatally mainly as a liver-derived endocrine hormone (Sjögren et al., 1999), the expression of *Igf1, Igf2*, and their receptor *Igf1r* in developing teeth (Joseph et al., 1994, **Table S4**) suggest a paracrine involvement of IGF signaling in tooth size regulation. Later during tooth development, IGF signaling is also important for tooth attachment to the jaw, as it regulates periodontal ligament formation (Jing et al., 2022).

**Figure 5.**
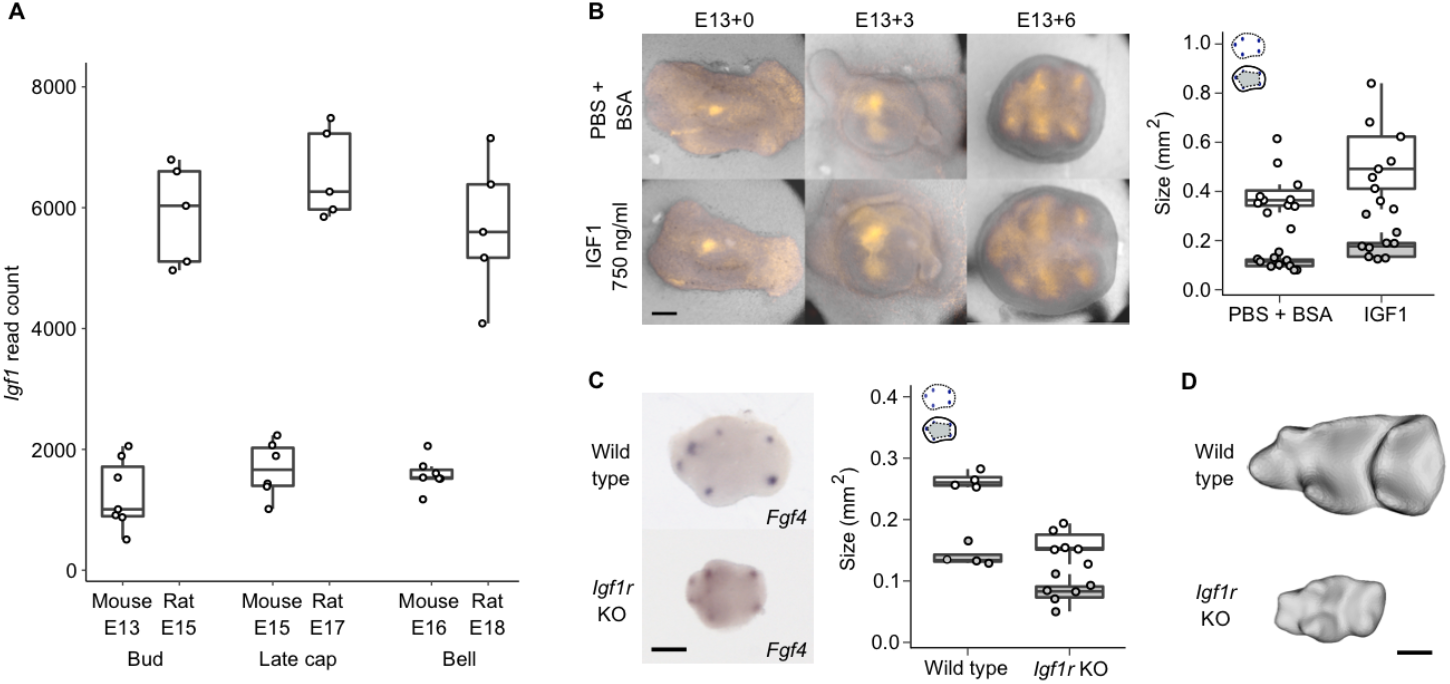
Modifying IGF1 signaling is sufficient to scale both tooth size and cusp patterning. **A**, *Igf1* is upregulated in the rat molars (*n* = 5 for all stages) compared to mouse molars (*n* = 7 per stage except late cap stage *n* = 6). **B**, IGF1-treated molars are larger (*n* = 9, *p* = 0.0339) and have more spread secondary enamel knots *ex vivo* (*n* = 9, *p* = 0.0001) than the controls (*n* = 11 for both measurements). **C**, *Igf1r* KO mouse molars are smaller (E18, *n* = 6, *p* < 0.0045) and their secondary enamel knots are less spread (*n* = 6, *p* < 0.0055) than those of wild type mouse molars (E17, *n* = 4). **D**, Wild type and *Igf1r* KO mouse molars at E19 when all the main cusps are visible. Boxes enclose 50% of observations, the horizontal bar denotes the median, and whiskers extend to last values within 1.5 interquartiles. Anterior to the left, buccal to the top. Scale bars, 200 µm.

To analyze the effects of IGF signaling on molar development experimentally, we first tested whether the IGF1 protein is capable of scaling up mouse molars *ex vivo*. IGF1 has been reported to increase tooth size in cultured or bioengineered molars (Young, 1995; Oyanagi et al., 2016), but its effects on normal patterning of enamel knots and cusps have not been studied. To visualize cusp patterning in culture, we used Fucci-red cell-cycle reporter mice and cultured their molars from bud stage (E13.5) with or without recombinant IGF1 protein (Methods). As the secondary enamel knots are non-proliferative, they become visible in Fucci-red mice before differentiation of the rest of the crown. We found that the size of IGF1-treated teeth is 1.35 times larger on average than that of the controls (**Fig. 5B**, *p* = 0.0339, **Table S5**). Similarly, the spacing of the secondary enamel knots of the treated teeth has increased in unison with the tooth size (**Fig. 5B**, *p* < 0.0001, **Table S5**), suggesting that excess IGF1 can both increase tooth size and scale the patterning so that the shape remains the same.

To examine how dependent tooth development is on canonical IGF signaling, we used an *in vivo* model of the *Igf1r* null mutant (*Igf1r*-KO) mouse (Holzenberger et al., 2003). Because these mice die perinatally, we analyzed the embryonic development of the molars (Methods). *Fgf4* expression in bell stage molars showed reduced spacing of the secondary enamel knots (*p* = 0.0055), as well as reduced tooth size, in the *Igf1r*-KO molars (**Fig. 5C**, *p* = 0.0045, **Table S6**), suggesting that the patterning process is downscaled in the mutant teeth. At E19, the *Igf1r*-KO molars have acquired all the main cusps even though they are only 0.33 in size compared to the corresponding wild type mouse molars (**Fig. 5D, Table S2**).

Taken together, IGF signaling appears to be sufficient to change both the size of the tooth and the patterning process, which in combination provides a mechanism for shape-invariant scaling. This inference raises the question of how IGF1 affects induction of secondary enamel knots, because the characteristic roles of IGF signaling are associated with growth and metabolism (Dupont & Holzenberger 2003; LeRoith et al., 2021), not patterning.

### IGF1 inhibits the expression of genes required for cusp patterning

To identify the downstream effects of IGF1 signaling in teeth, we treated cap stage (E14) mouse lower molars with recombinant IGF1 protein for 6 hours (Methods) followed by RNAseq analysis of differential gene expression. The IGF1 treatment shows the expected bias towards upregulation of metabolic and biosynthesis related genes (**Fig. S2A**), whereas the downregulated genes appear to be related to developmental regulation (**Fig. S2B**). A closer examination of these results shows that several DNA replication markers were upregulated by IGF1 **(Fig. 6A)**, implicating stimulation of cell proliferation. In strong contrast, there was a total lack of upregulation of any of the known developmental genes (Hallikas et al., 2021) required for normal tooth morphogenesis (**Fig. 6B, Table S7**). Instead, we found eight tooth genes to be downregulated (*p* < 0.05 **Fig. 6B, Table S7**), five of which are expressed in the enamel knots. Of the downregulated genes, *Lef1* and *Bmp4* are required for the induction of the molar enamel knots (Sasaki et al., 2005; Jia et al., 2013), while the others alter cusp patterns when mutated (Hallikas et al., 2021). Overall, IGF signaling appears to have a dual role in tooth development: induction of growth and, at the same time, inhibition of enamel knot driven cusp patterning. This result may also help to explain why attempts to increase tooth size experimentally by increasing tissue size or by recombining tissues lead to an increase in cusp number (Cai et al., 2007; Ishida et al., 2011).

**Figure 6.**
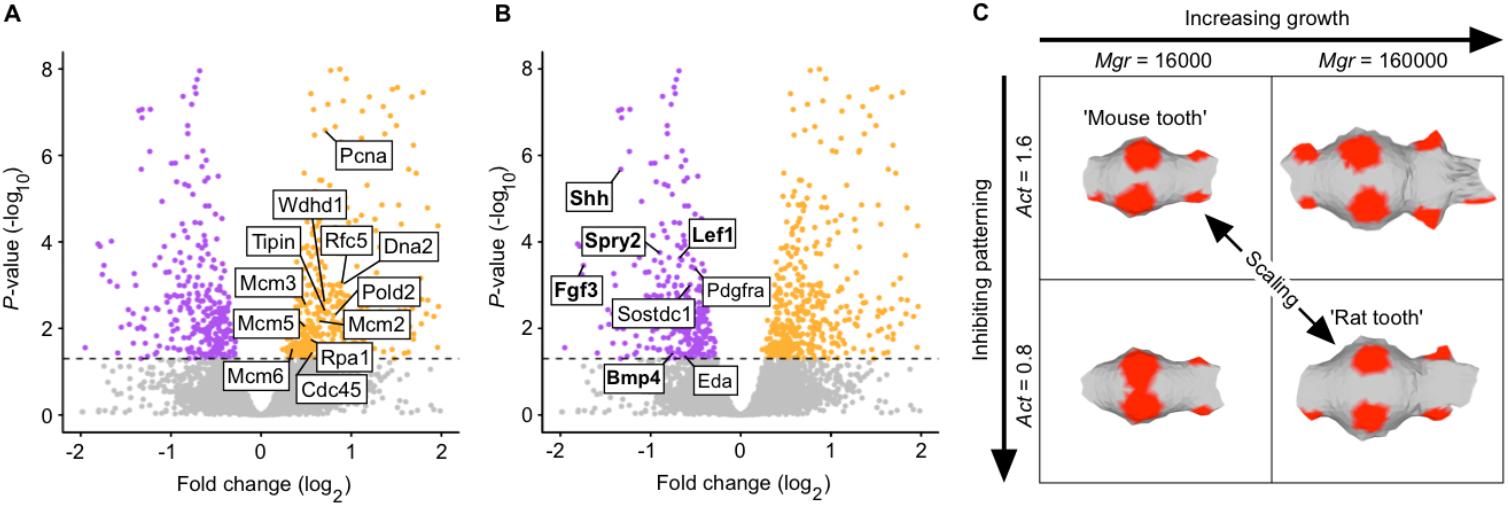
Simultaneous promotion of overall growth and inhibition of cusp patterning by IGF1 provides a mechanism to scale teeth. **A**, Mouse molars treated with IGF1 protein for six hours show upregulation (orange) of 12 DNA replication markers (GO:0006260), *n* = 5 for both the treatments and controls. **B**, In contrast to growth promoting effects (**A**), genes required for normal tooth development show only downregulation in IGF1-treated molars (downregulated genes in purple). Enamel knot expressed genes are in bold. Horizontal line denotes padj = 0.05. Volcano plots are zoomed to the genes of interest, see **Fig. S2** for the overall fold enrichment of GO categories. **C**, Computer simulations of molar development using ToothMaker show that changing proliferation rate (*Mgr*) or activator autoregulation (*Act*) alone changes the pattern (forming secondary enamel knot regions shown in red). By increasing growth and by decreasing activation, which mimics the effects in (**A**) and (**B**), the simulated mouse tooth can be scaled up. See text and Methods for details.

To further investigate the principle of dual requirement of growth and patterning regulating scaling, we used a computational model of tooth development to scale teeth (ToothMaker; ref. Harjunmaa et al., 2014). This morphodynamic model integrates signaling and tissue growth to simulate tooth development, and it has been used in experimental and evolutionary studies (Harjunmaa et al., 2014, Renvoise et al., 2017; Savriama et al., 2018; Couzens et al., 2021; Thiery et al., 2022), but not to examine scaling. As a starting point, we used the simulated mouse molar from previous studies (Harjunmaa et al., 2014, Renvoise et al., 2017) and increased its size (Methods). Increasing only the growth resulted in additional secondary enamel knots and altered cusp pattern (**Fig. 6C, Table S8**). However, by simultaneously decreasing the activator required for enamel knot induction, we obtained a larger tooth that retains the mouse pattern with more widely spread enamel knots (**Fig. 6C, Table S8**). Decreasing activation resulted in the requirement of a larger number of activator producing cells, hence larger size, to reach the threshold to induce the secondary enamel knots. Taken together, we interpret these results to support the role of IGF signaling, likely through changes in many of the pathway genes (**Table S4**), as a single ‘dial’ that simultaneously promotes growth and inhibits patterning.

### Comparative data on mammalian teeth support universality of the scaling mechanism

Because our inferences on the scaling of patterning were based on two murine species, we wanted to examine a broader range of species and sizes. Here we took advantage of our observation that differences in tooth width between the mouse and rat appear to become discernable relatively late, beginning with the cap stage (**Fig. 3A, Table S1, S2**). Frontal histological sections of cap-stage teeth are available for different species in the literature as also in museum collections, providing data to use tooth width as a proxy for tooth size (**Fig. 7A**, Methods, **Table S9**). We therefore measured early cap-stage widths from developing molars and corresponding fully formed tooth widths of 14 mammalian species, ranging in size from the shrew (*Sorex araneus*) to the elephant (*Loxodonta africana*) (**Table S9**). The measurements show that even though these teeth vary over 74-fold in final, mineralized width, the early cap-stage widths exhibit very little change in size (**Fig. 7B, Table S10**). This means that whereas the fully formed teeth scale with body size, the early cap stage tooth germs are relatively size-invariant.

**Figure 7.**
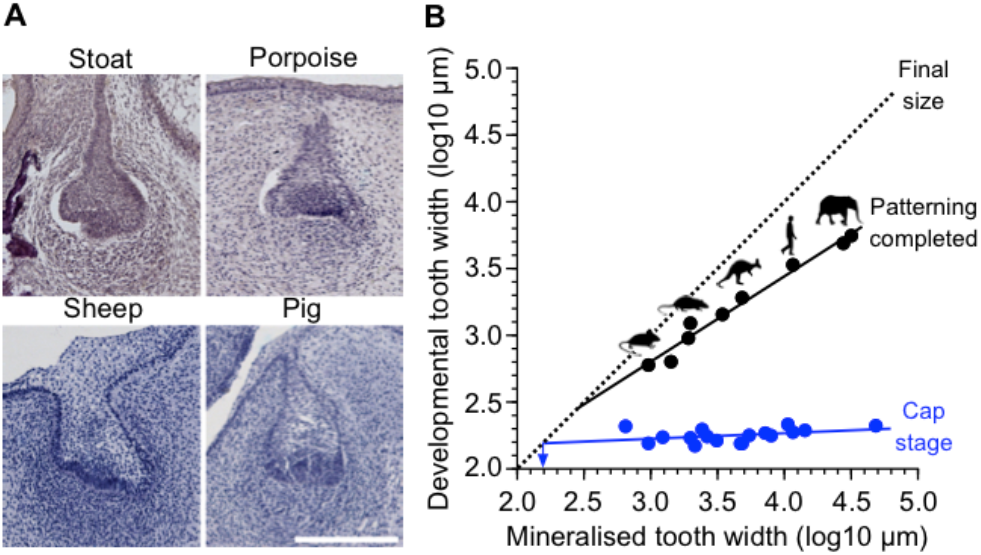
Patterning scales across mammals. **A**, Frontal sections of developing teeth of various mammals show similar bucco-lingual widths of tooth germs at early cap stage. The sections show dp4 (sheep), p4 (stoat), and dp3 (pig). The porpoise tooth identity cannot be determined. **B**, The cap stage widths (blue) do not show a marked increase with the final mineralized tooth widths (regression slope is 0.043 and the intercept is 2.095, *r*^2^ = 0.163), and the regression-line extrapolated minimum tooth width is 154 µm for single cusped teeth (arrow). In contrast, the widths when the patterning is completed increase as the teeth become larger (black line, regression slope is 0.640 and the intercept is 0.885, *r*^*2*^ = 0.976). The point when tooth development reaches the final tooth size is marked with the dashed diagonal (x-and y-axis values are the same). For sample details, see **Table S9**. Scale bar, 200 µm in (**A**).

The fully formed, final tooth size can be expected to be close to the cap-stage width in single cusped teeth when their cervical loops grow directly downwards. The average width of our cap-stage measurements was 178 µm (**Table S9**). Taking into account the regression slope, the extrapolated minimum width for the cap stage would be even smaller at 154 µm, suggesting this as the theoretical lower limit for tooth size in mammals (arrow at 2.2 log10 in **Fig. 7B, Table S10**). Notwithstanding that this limit should be considered an approximation (**Table S10**), it is still instructive to consider the empirical data. Mammalian teeth can be less than 500 µm in diameter, even with multiple cusps such as the mouse third molars. Experimentally, extreme reduction in tooth size has been achieved in mice with activated epithelial Wnt signaling (Järvinen et al., 2006). In these mice, teeth are continuously generated, typically with round, peg-like morphology (Järvinen et al., 2006). As the size distribution of these teeth has not been examined previously, we quantified the sizes of 42 mineralised teeth obtained from a single molar germ transplant experiment, cultured under the kidney capsule (Järvinen et al., 2006). The frequency distribution of the teeth shows (**Fig. S3**) that, towards the smaller teeth, their size distribution falls steeply around 200 µm, with only one tooth being clearly narrower than the predicted minimum (98 versus 154 µm). Moreover, teeth with two well-differentiated cusps appear to be at least about 400 µm wide (**Fig. S3**).

Although the data to compare scaling of patterning are more limited, we nonetheless obtained tooth widths for seven species at the stages when the last forming cusps have just been initiated during ontogeny (Methods, **Table S9**). Unlike the early cap-stage tooth germs, these fully patterned teeth scale with the final tooth width (**Fig. 7B**). Considering again the widths of developing teeth, the best-fit line for cap stage teeth extrapolated towards zero overlaps with a theoretical minimum at 286 µm (2.5 log10 in **Fig. 7B, Table S10**). This indicates that patterning is truncated in smaller teeth and agrees with the lack of teeth with second cusps in the transplant experiment (**Fig. S3**). To the extent that these values are representative of mammalian tooth development in general, the downscaling capacity of teeth appears to be progressively constrained when teeth become less than half a millimeter in diameter.

For larger teeth, the consequence of the size-invariant initiation, followed by the scaling of the patterning, is a progressive increase of growth during patterning as the teeth become larger. For example, whereas linear size in the mouse increases by 3.8 times in the cap stage relative to the end of the patterning, the comparable increase is 15.4 times in the human (calculated for 1 mm and 10 mm sized teeth, respectively). After patterning, the final increases in tooth sizes are 1.6 and 3.6 times for the mouse and human, which are only about 0.4 and 0.2 times the comparable increases during patterning, respectively. In other words, patterning of larger teeth encompasses an increasingly large share of the cell divisions needed to reach the final size (**Fig. 7B**).

## DISCUSSION

Evolution of tooth size has had a central role in the reconstruction of body size evolution in mammals (Damuth & MacFadden, 1990; Gingerich, 1982; Damuth & MacFadden, 1990; Alroy, 1998; Smith et al., 2010; Smith et al., 2016; D’Ambrosia et al., 2017). Also, as size affects many aspects of an animal’s ecology, size changes alone are often used as a diagnostic feature to delineate species. Evolutionary changes in body size have been frequent in mammalian evolution, and tooth size seems to track these changes closely, although with a slight delay when the change is very fast (e.g. domesticated mammals, see Clauss et al., 2022). Here we investigated how the scaling of teeth can be achieved during development. Comparisons of mouse and rat molars show that scaling is already active during the patterning phase of tooth development. Tooth patterning, which is responsible for the formation of species-specific cusp patterns, is a critical period of morphogenesis that is sensitive to mutations in many regulatory genes (Hallikas et al., 2021). Our experimental data and modeling results implicate IGF signaling as a mechanism for scaling both the patterning and the size. This includes the well-established role of IGF signaling in promoting growth (Dupont & Holzenberger, 2003; LeRoith et al., 2021), and also the regulation of secondary enamel knots by inhibiting their activation (**Figs 5, 6**). This in turn would result in the requirement for a larger number of cells, and larger size, to reach the signaling threshold for cusp formation in larger teeth (**Fig. 6C**). More generally, our results further underscore the diverse roles that IGF signaling appears to play in developing teeth (Koffi et al., 2021; Jing et al., 2022).

One obvious question that arises from these analyses is why should scaling and patterning be integrated. One possible answer is that larger teeth retain patterning control for a progressively larger share of their increase in size (**Fig. 7B**), which in turn may minimize accumulation of harmful changes in shape caused by growth alone. Upscaling the patterning may also have the side effect of allowing large teeth to elaborate cusp patterns for longer periods of developmental time. Cursory analyses of dental diversity have shown that larger teeth tend to have more cusps (Jernvall, 1995), and this could be in part due to the scaling of patterning. Another contributing factor in the increase of cusp number in larger teeth is the prevalence of herbivory in large mammals. Herbivores have relatively complex teeth (Evans et al., 2007), which are presumably easier to achieve as the teeth become larger.

The predicted minimum tooth sizes with single and additional cusps (**Fig. S3, Tables S9, S10**) may be relatively close to some of the teeth in early mammaliforms (Luo, 2007; Gill et al., 2014). Making even smaller teeth might require smaller cell size, or alternative mechanisms for patterning (Larionova et al., 2021). Small multicusped teeth do occur in reptiles (Lafuma et al., 2021) and fish (Streelman et al., 2003); at least in sharks tooth cusp patterning has been proposed to be relatively mammal-like (Thiery et al., 2022).

The mammalian dentition evolved from a common ancestor with relatively simple teeth lacking lateral cusps (e.g., Gill et al., 2014). The subsequent lateral expansion can be considered an evolutionary novelty and a prerequisite for the acquisition of tribosphenic molars which combine slicing and crushing functions (Luo, 2007; Gill et al., 2014; Couzens et al., 2021). The sequence of evolutionary changes leading to tribospheny may explain the relatively late onset of lateral expansion of molars during development (**Fig. 2, 3, Table S1, S2**). This stepwise development also enables the continuing differentiation of dentitions into laterally expanded molars and laterally narrow anterior teeth, and suggests that shape-invariant scaling of teeth is not the developmental default but an actively retained scaling relationship involving all the steps of morphogenesis.

Because organs generally scale with body size, we predict that comparable scaling of patterning, possibly driven by IGF signaling, may occur in most organs. At least in the case of teeth, which have determinate growth, the final tooth size can be used to predict the size of patterning phase during development. Thus, in addition to being useful in inferring body size, tooth size is also informative about development.

## MATERIALS AND METHODS

### Animals

All mouse and rat studies were approved and carried out in accordance with the guidelines of the Finnish national animal experimentation board under licenses KEK16-021, KEK19-019 (mice) and KEK17-026, KEK14-026 and KEK13-014 (rats).

We used wild type outbred NMRI mice and RccHan:Wist Wistar rats for micro-computed tomography and *in situ* hybridization and inbred C57BL/6JOlaHsd mice and DA/HanRj rats for transcriptomics. Tissue culture experiments were carried out using Fucci mouse line expressing nuclear red (mKO-Cdt1) in G1 cell cycle phase in NMRI background (Sakaue-Sawano et al., 2008). IGF1R-KO mice (Holzenberger et al., 2000, 2003) were kept in outbred 129S2/SvPasCrl background. Embryo age was determined based on vaginal plug appearance (embryonic day, E0). We confirmed the comparable dental stage by comparison of tooth morphology and the appearance of dental signaling centers.

### Micro-computed tomography (µCT)

Mandibles of E13-E17 mouse and E15-20 rat embryos were fixed overnight in 4% paraformaldehyde, dehydrated gradually to 70% ethanol and stored at +4 °C. Postnatal mandibles were fixed in 4% PFA for 1-2 days (depending on size) and gradually dehydrated to 70% ethanol. Phosphotungstic acid (PTA) was used to increase soft-tissue contrast for µCT imaging (Metscher, 2009). Samples were stained in 0.3% PTA (Sigma Aldrich) in 70% ethanol for 48-72 hours at +4 °C and stored in 70% ethanol. For scanning, samples were embedded in 1% low-melting point agarose dissolved in MilliQ water. Scanning was carried out using Bruker 1272 µCT scanner with polychromatic cone beam X-ray source (Hamamatsu L11871 20, 20-100 kV), 11-Megapixel xiRAY X-ray CCD camera with Onsemi KAI-11002 sensor fiber-optically coupled to P43 scintillator. Embryonic samples were scanned using 0.25 mm aluminum filter at 60 kV and 166 µA. Postnatal samples were scanned using 0.5 mm aluminum filter at 70 kV and 142 µA. The voxel size used varied between 1-4 µm depending on the specimen size. Reconstruction was carried out using Bruker NRecon software (version 1.6.10.1), and ring artefact correction was used when necessary. Scanning of two *Loxodonta* fetuses (University Museum of Zoology Cambridge or UMZC 2011.10.1 and UMZC 2013.7) followed PTA staining protocols in Table 2 of (Metscher, 2009). Both specimens were scanned at the Cambridge Biotomography Centre on a Nikon-Xtek H-225-ST. UMZC 2013.7 was in 0.3% PTA solution for 8 weeks and scanned using 0.5 to 1 mm copper filters at 140-142 kV and 240-340 µA; UMZC 2011.10.1 was in 0.3% PTA for 1 week and scanned without a filter at 110 kV and 167 µA.

### Segmentation and tooth measurements

Segmentation of molars was carried out using Avizo (release 9.0.1). The epithelium was segmented manually using lasso tool, and the mesenchyme using brush tool. Every 3^rd^ to 5^th^ section was drawn and the sections in between interpolated, but the accuracy of automatic interpolation was confirmed manually in each section, and corrected when necessary. After segmentation, the binary stack was opened in Fiji (Schindelin et al., 2012) and smoothed using Gaussian blur 3D-tool with x, y and z sigma of 3. A standard deviation Z-project was taken from occlusal side and the tooth was measured using magic wand (area) and bounding box (maximum antero-posterior length and bucco-lingual width). Logistic curve fitting was done with PAST (Hammer et al., 2001). We report the results using two-dimensional areas because they are commonly used in evolutionary analyses, and because these were obtainable for both *in vivo* and *ex vivo* data. For measurements from histological section, the bucco-lingual widths of teeth of different species were acquired from the Museum of Natural History Berlin, Germany. Histological slides were imaged using Zeiss Axioskop, Plan-Neofluar 5x objective and Leica DFC490 camera. Additional measurements were done from the literature (**Table S9**).

### Orientation patch count

For OPC measurements the segmented mesenchymes were saved as .stl surfaces using Fiji 3D viewer. The surfaces were handled in Meshlab (version 2021.10). The faces were inverted and the basal surface of the mesenchyme was removed using the Z-painting tool to limit the analysis only to the occlusal surface. Teeth were oriented and the scan resolution differences were corrected for by dividing the original face number with ((4/x)^2) where x is the original resolution of the scan in micrometers. The acquired value was used as target number of faces in quadric edge collapse decimation tool. The surfaces were smoothed with 50 steps using Laplacian smooth to remove segmentation artifacts and to focus on the overall surface topography, and simplified to 4000 faces each using quadric edge collapse decimation with planar simplification weight set to one, in order to produce relatively uniform distribution of triangles. Although the use of a similar face count is used to remove the effect of size, we note that this procedure still results in smaller triangles in teeth with low relief. In our data this does not affect the pattern of results because the relief increase similarly between the species. OPC values (OPCR) of resolution-corrected surfaces were acquired using Morphotester (version 11.2, Winchester, 2016) with a minimum patch count of 6 (roughly matching 3 pixels in raster based OPC). Visualization was done with modified R script in molarR (Pampush et al., 2016).

### Probe synthesis

For interspecies comparison, species-specific probes were designed for each marker. Probes were designed to bind the same part of the mRNA in each species. Species-specific primers used for preparing probes are listed below. cDNA was prepared from mouse and rat embryonic molar tooth RNA (extracted using RNeasy Plus Micro kit, Qiagen, Düsseldorf, Germany). cDNA constructs were inserted in TOPO II PCR-plasmids and using TOPO TA Cloning kit with chemically competent cells according to manufacturer’s protocol (Thermo Fisher Scientific, Waltham, Massachusetts, U.S.). Prior to *in vitro* RNA synthesis, plasmids were extracted using Miniprep kit (Qiagen, Düsseldorf, Germany). Plasmids were linearized and probes were prepared as described in Wilkinson & Nieto (1993) using digoxigenin-conjugated nucleotides (Roche, Basel, Switzerland). Sense probes were used to confirm specificity of the antisense probes. The following primers were used (forward and reverse primers are listed respectively): CGTAAGTCCTTCACCAGCTTG and GCTGACCCCTTTAGCCTACA for mouse *Shh*, CTTAGATCCTTCACTAACTTGGTG and GCTGACCCCTTTAGCCTACA for rat *Shh*, GGAAGGGTAATTACTGGACTC and ATGAGGCTGTTGACCATGCTG for mouse *Foxi3* (Ohyama & Groves, 2004), GAAAAGGTAATTACTGGACTC and ATGAGGCTGTTGACCATGCTG for rat *Foxi3*, CAACGTGGGCATCGGATTC and CCTCATGGTAGGCGACACT for mouse *Fgf4*, AGGCTGCGGAGACTCTACTG and GAAACTCGGTTCCCCTTCTT for rat *Fgf4*.

### Whole mount *in situ* hybridization

Mandibles of E12-E14.5 mouse embryos and E14-E17 rat embryos were dissected for placode and primary enamel knot analysis. To be able to detect the secondary enamel knots, E16-17 mouse molars and E18-20 rat molars were separated from the mandible and the thick outer enamel epithelium was removed. All samples were fixed overnight in 4% paraformaldehyde, dehydrated to 100% methanol and stored at -20 °C. A routine *in situ* hybridization protocol (Wilkinson & Nieto, 1993) was used with the following alterations: hydrogen peroxide and glutaraldehyde were not used, proteinase K (Roche, Basel, Switzerland) was used in 7 mg/ml concentration, before prehybridization samples were treated with acetic anhydride in 0.1 M triethanolamine for 10 minutes, hybridization buffer had additional 50 µg/ml yeast tRNA and 1x Denhardt’s solution (Invitrogen, Waltham, Massachusetts. U.S.), all post-hybridization washes were carried out using 5x SSC, 50% formamide, 0.1% Tween20, blocking and antibody solutions had 1% Boehringer’s blocking reagent (Roche, Basel, Switzerland), and 10% and 1% of goat serum, respectively. Alkaline-phosphatase bound anti-digoxigenin antibody (11093274910, Roche, Basel, Switzerland) was used to detect the mRNA probe. Levamisole was not used in alkaline phosphatase buffer, and BM-purple (Roche, Basel, Switzerland) was used as alkaline phosphatase substrate. Samples were imaged using Zeiss Lumar V12 stereo microscope, Apolumar S 1.2x objective and AxiocamICc1 camera.

### Placode and signaling center measurements

The placode area, the initiation knot area and the primary enamel knot area were measured using Fiji (Schindelin et al., 2019). Samples with weak staining were excluded. When both right and left sides of the jaw were available, left side was used. The images were converted to 8-bit and pixels included in the expression area were defined as: (tissue median pixel value – expression area minimum pixel value)/2 + expression area minimum pixel value. The area enclosed by the secondary enamel knots was determined by drawing a polygon between enamel knot centers using the polygon tool in Fiji. Only teeth where at least five enamel knots were present, but without distinct development of the cusps, were measured. The tooth areas were measured using polygon tool in Fiji. Randomization test with 10,000 permutations in R (modified from Kuiper & Sklar, 2012) was used to test differences between samples for the placode size, initiation knot size, primary enamel knot size, the area enclosed by the secondary enamel knots, and tooth size during patterning (**Figs 2, 4, 5B, C**). All the *p*-values are reported as two-tailed. Alternative methods to threshold the expression domains do not alter the pattern of results. Mouse strains used in controls were the same as their experimental contrasts.

### Cell size measurements

Cell sizes of mouse and rat embryonic molars were determined by staining 6 µm thick histological sections with DiI (Thermo Fisher Scientific, Waltham, Massachusetts, U.S.) and Hoechst nuclear stain (Invitrogen, Waltham, Massachusetts. U.S.). Sections were rehydrated gradually to RO H_2_O, washed in PBS + 0.3% Triton-X, and incubated in DiI (25 mg/ml in absolute ethanol stock dissolved in PBS in 1:200 ratio) for 45 minutes. Sections were washed in PBS (4x5 min) and incubated in 1:2000 Hoechst for two hours prior to mounting. Sections were imaged using Zeiss Axio Imager.M2, Axiocam HRc camera with Zeiss 40x Plan Neofluar objective. Cell perimeters were measured using lasso tool in Fiji.

### Tooth cultures

E13 mouse molars were dissected and cultured at 37 °C with 5% CO_2_ using a Trowell type organ culture as described previously (Närhi & Thesleff, 2010). Media was supplemented with ascorbic acid (100 µg/ml, Sigma-Aldrich, Burlington, Massachusetts, U.S.) and 750 ng/ml recombinant mouse IGF1 protein (791-MG-050, Bio-Techne, Minneapolis, Minnesota, U.S.) in 1x PBS + 0.1% BSA or similar volume of PBS + 0.1% BSA in controls. Samples were imaged daily using Zeiss Lumar V12 stereo microscope, Apolumar S 1.2x objective and AxiocamICc1 camera. A drop of media (7 µl) was added on top of each sample daily to prevent the samples from drying. Media was changed every other day. The cultures were stopped when 5-6 secondary enamel knots were visible. The distribution of the secondary enamel knots was defined by drawing a polygon between enamel knot centers using the polygon tool in Fiji. The tooth areas were measured using polygon tool in Fiji.

### IGF1 induction

E14 mouse molars were dissected and cultured in a hanging drop culture (Närhi & Thesleff, 2010) for 6 hours pairwise so that from each embryo one tooth was treated with control media and one with IGF1-containing media (media constituents and concentrations described in the previous section). Right and left sides were balanced, *n* = 5 for both the treatments and controls.

### Transcriptomics

Wild type tooth germs were dissected from E13, E15 and E16 mouse molars. Teeth of corresponding morphological stages were dissected from E15, E17 and E18 rats. Minimal amount of surrounding tissue was left around the tooth germ, at the same time making sure that the tooth was not damaged in the process. The tissue was immediately stored in RNAlater (Qiagen, Düsseldorf, Germany) at −80 °C for RNAseq. For RNAseq, each tooth was handled individually. Seven biological replicates were collected for mouse and five biological replicates for rat. Numbers of left and right teeth were balanced. The samples of IGF1 induction experiment were processed similarly. Samples were homogenized in TRI Reagent (Merck) using Precellys 24-homogenizer (Bertin Instruments). RNA was extracted by guanidium thiocyanate-phenol-chloroform method and purified using RNeasy Plus micro kit (Qiagen GmbH). The RNA quality of representative samples was confirmed using 2100 Bioanalyzer (Agilent). The purity of RNA was analyzed using Nanodrop (Thermo Fisher Scientific). RNA concentration was measured by Qubit 3.0 (Thermo Fisher Scientific). The complementary DNA (cDNA) libraries were prepared using Ovation Mouse RNAseq System and Ovation Rat RNAseq System (Tecan). Gene expression levels were measured using RNAseq (platforms GPL19057, Illumina NextSeq 500). The RNAseq reads of mouse and rat were evaluated and bad reads were filtered out using FastQC (version 0.11.8, Andrews et al., <u>2012</u>), AfterQC (version 0.9.6, Chen et al., 2017) and Trimmomatic (version 0.39, Bolger et al., 2014), and ribosomal RNA was removed using Sortmerna (Kopylova et al., 2012). The number of remaining, good reads varied between 30M and 90M in the rat samples and 40M and 65M in the mouse samples, and 8.9M and 22.7M reads for IGF1-induction experiment. Mouse and rat reads were aligned using Salmon (version 0.99.0, Patro et al., 2017) to GRCm38 (Ensembl release 100) cDNA and Rnor_6.0 (Ensembl release 99) cDNA, respectively. For mouse and rat comparison, 16,604 one-to-one orthologous genes were found between mouse and rat using Ensembl Biomart tool (version 2.50.3, Kinsella et al., 2011). 126 additional one-to-one orthologues were added using Inparanoid8 (Sonnhammer & Östlund, 2015) in which gene pairs with bootstrap scores of 100% were selected. Only these one-to-one-orthologues found with Biomart and Inparanoid8 (version 8.0, data downloaded on June 2020) were used for further analysis. The mouse and rat output transcript ID’s of Salmon were converted to mouse gene ID’s using Ensembldb (Rainer et al., 2019) and Tximport (version 1.22.0, Soneson et al., 2015), allowing comparison of mouse and rat read counts. Deseq2 (version 1.34.0, Love et al., 2014) was used to normalise the read counts by library size and composition as well as transcript length. For Gene Ontology (GO) term analyses of biological processes, PANTHER 17.0 (Thomas et al., 2022) was used to examine up-and downregulated genes of the IGF1-induction experiment. Fold enrichment analysis was done using PANTHER overrepresentation test (Release 20221013, GO Ontology database DOI: 10.5281/zenodo.6799722 Released 2022-07-01) with default Fisher’s exact test and False Discovery Rate correction (Huaiyu et al., 2019). All transcriptome data are available in GEO at https://www.ncbi.nlm.nih.gov/geo/query/acc.cgi, reference numbers GSE142199, GSE158697, and GSE218338.

### Computational modeling

ToothMaker (Harjunmaa et al., 2014) was used to investigate the scaling of mouse molar simulations used in previous studies (Harjunmaa et al., 2014; Renvoise et al., 2017, **Table S8**). The model implements experimentally inferred genetic interactions with tissue biomechanics to simulate tooth development. The logic of the model is morphodynamic (Salazar-Ciudad & Jernvall, 2010) in that signaling regulating patterning happens concomitantly with growth. Starting from parameters used previously to simulate mouse molar development (Harjunmaa et al., 2014; Renvoise et al., 2017), we systematically increased the mesenchymal proliferation rate (*Mgr*) and decreased the auto-activation of activator (*Act*). These changes simulate increases in growth and inhibition of patterning, respectively (**Table S8)**. All simulations were run for the same number of iterations (14,000), which cover the development up to early bell stage (approximately five days after the placode stage). ToothMaker is available at https://github.com/jernvall-lab/ToothMaker.

## Acknowledgements

We thank M. Fortelius, V. Hietakangas, J. Laakkonen, M. Mikkola, I. Salazar-Ciudad, K. Kavanagh, and members of Jernvall lab for comments and discussions on this work; N. Di-Poï and J. Laakkonen for help with comparative material; H. Suhonen for help with microCT imaging; A. Viherä, R. Savolainen, R. Murray, M. Mäkinen, O. Saarnisalo, and M. G. Varghese for technical assistance; P. Auvinen, L. Paulin and P. Laamanen at DNA Sequencing and Genomics Laboratory; RIKEN BioResource Center through the National Bio-Resource Project of the MEXT, Ibaraki, Japan providing the Fucci mice; P. Giere (Museum für Naturkunde, Berlin) and J. Granroth (Finnish Museum of Natural History, Helsinki) for access and help with museum collections. For access to and assistance with *Loxodonta* specimens, R.J.A. thanks F. Stansfield, L. Hautier, and the late R. T. Allen. This study was supported by the Academy of Finland, Sigrid Jusélius Foundation, and John Templeton Foundation (J.J.), Doctoral Programme in Biomedicine (M.M.C.), and by NIDCR R01-DE027620 and R35-DE026602 (T.J.H. and O.D.K.).

## Author contributions

M.M.C. and J.J. conceived the project. M.M.C. and V.V. obtained and M.M.C., O.E.S. and J.J. analyzed the phenotypic data. M.M.C. performed culturing experiments and measurements. M.M.C., O.H., and R.D.R. performed transcriptomics. T.J.H. developed computational tools. M.M.C. and J.J. compiled the comparative data. J.-C.F., M.H., R.J.A., and O.D.K. contributed materials, observations and scientific interpretations. M.M.C. and J.J. integrated the analyses and wrote the paper with input from all authors.

## REFERENCES

Alroy, J. (1998). Cope’s Rule and the Dynamics of Body Mass Evolution in North American Fossil Mammals. Science, 280(5364), 731–734. doi: 10.1126/science.280.5364.731

Alvesalo, L., & Tigerstedt, P. M. A. (1974). Heritabilities of human tooth dimensions. Hereditas, 77, 311–318. doi: 10.1111/j.1601-5223.1974.tb00943.x

Andrews, S. (2010). FastQC: A Quality Control Tool for High Throughput Sequence Data [Online]. Available online at: http://www.bioinformatics.babraham.ac.uk/projects/fastqc/

Bolger, A. M., Lohse, M., & Usadel, B. (2014). Trimmomatic: a flexible trimmer for Illumina sequence data. Bioinformatics, 30(15), 2114–2120. doi: 10.1093/bioinformatics/btu170

Boulan, L., & Léopold, P. (2021). What determines organ size during development and regeneration? Development, 148(1), 1–9. doi: 10.1242/dev.196063

Butler, P. M. (1967). The prenatal development of the human first upper permanent molar. Archives of Oral Biology, 12(4), 551–563. doi: 10.1016/0003-9969(67)90030-1

Cai, J., Cho, S.-W., Kim, J.-Y., Lee, M.-J., Cha, Y.-G., & Jung, H.-S. (2007). Patterning the size and number of tooth and its cusps. Developmental Biology, 304(2), 499–507. doi: 10.1016/j.ydbio.2007.01.002

Chen, S., Huang, T., Zhou, Y., Han, Y., Xu, M., & Gu, J. (2017). AfterQC: automatic filtering, trimming, error removing and quality control for fastq data. BMC Bioinformatics, 18(Suppl 3), 80. doi: 10.1186/s12859-017-1469-3

Clauss, M., Heck, L., Veitschegger, K., & Geiger, M. (2022). Teeth out of proportion: Smaller horse and cattle breeds have comparatively larger teeth. Journal of Experimental Zoology Part B: Molecular and Developmental Evolution. doi: 10.1002/jez.b.23128

Copes, L. E., & Schwartz, G. T. (2010). The scale of it all: postcanine tooth size, the taxon-level effect, and the universality of Gould’s scaling law. Paleobiology, 36(2), 188–203. doi: 10.1666/08089.1

Couzens, A. M. C., Sears, K. E., & Rücklin, M. (2021). Developmental influence on evolutionary rates and the origin of placental mammal tooth complexity. Proceedings of the National Academy of Sciences, 118(23), e2019294118. doi: 10.1073/pnas.2019294118

Damuth J. & MacFadden B. J. (1990). Body size in mammalian paleobiology: Estimation and biological implications. Cambridge University Press.

D’Ambrosia, A., Clyde, W. C., Fricke, H. C., Gingerich, P. D. & Abels, H. A. (2017). Repetitive mammalian dwarfing during ancient greenhouse warming events. Science Advances, 3, doi: 10.1126/sciadv.1601430

Dupont, J., & Holzenberger, M. (2003). Biology of insulin-like growth factors in development. Birth Defects Research Part C: Embryo Today: Reviews, 69(4), 257–271. doi: 10.1002/bdrc.10022

Evans, A. R., Wilson, G. P., Fortelius, M., & Jernvall, J. (2007). High-level similarity of dentitions in carnivorans and rodents. Nature, 445(7123), 78–81. doi: 10.1038/nature05433

Gaunt, W. A. (1959). The development of the deciduous cheek teeth of the cat. Acta Anatomica, 38, 187–212. doi: 10.1159/000141527

Gaunt, W. A. (1961). The development of the molar pattern of the golden hamster (Mesocricetus auratus W.), together with a re-assessment of the molar pattern of the mouse (Mus musculus). Acta Anatomica, 45, 219–251. doi: 10.1159/000141753

Gill, P. G., Purnell, M. A., Crumpton, N., Brown, K. R., Gostling, N. J., Stampanoni, M., & Rayfield, E. J. (2014). Dietary specializations and diversity in feeding ecology of the earliest stem mammals. Nature, 512(7514), 303–305. doi: 10.1038/nature13622

Gingerich, P. D., Smith, B. H., & Rosenberg, K. (1982). Allometric scaling in the dentition of primates and prediction of body weight from tooth size in fossils. American Journal of Physical Anthropology, 58(1), 81–100. doi: 10.1002/ajpa.1330580110

Hallikas, O., Roy, R. D., Christensen, M. M., Renvoisé, E., Sulic, A., & Jernvall, J. (2021). System-level analyses of keystone genes required for mammalian tooth development. Journal of Experimental Zoology Part B: Molecular and Developmental Evolution, 336(1), 7–17. doi: 10.1002/jez.b.23009

Hammer, Ø., Harper, D.A.T., Ryan, P.D. (2001). PAST: Paleontological statistics software package for education and data analysis. Palaeontologia Electronica 4(1): 9pp.

Harjunmaa, E., Seidel, K., Häkkinen, T., Renvoisé, E., Corfe, I. J., Kallonen, A., … Jernvall, J. (2014). Replaying evolutionary transitions from the dental fossil record. Nature, 512(7512), 44–48. doi: 10.1038/nature13613

Harmansa, S., & Lecuit, T. (2021). Forward and feedback control mechanisms of developmental tissue growth. Cells & Development, 168, 203750. doi: 10.1016/j.cdev.2021.203750

Holzenberger, M., Leneuve, P., Hamard, G., Ducos, B., Perin, L., Binoux, M., & Bouc, Y. L. (2000). A Targeted Partial Invalidation of the Insulin-Like Growth Factor I Receptor Gene in Mice Causes a Postnatal Growth Deficit. Endocrinology, 141(7), 2557–2566. doi: 10.1210/endo.141.7.7550

Holzenberger, M., Dupont, J., Ducos, B., Leneuve, P., Géloën, A., Even, P. C., … Bouc, Y. L. (2003). IGF-1 receptor regulates lifespan and resistance to oxidative stress in mice. Nature, 421(6919), 182–187. doi: 10.1038/nature01298

Hovorakova, M., Lesot, H., Peterka, M., & Peterkova, R. (2005). The developmental relationship between the deciduous dentition and the oral vestibule in human embryos. Anatomy and Embryology, 209(4), 303–313. doi: 10.1007/s00429-004-0441-y

Hovorakova, M., Lesot, H., Peterka, M., & Peterkova, R. (2018). Early development of the human dentition revisited. Journal of Anatomy, 233(2), 135–145. doi: 10.1111/joa.12825

Hopkins, S.S.B. (2018). Estimation of Body Size in Fossil Mammals. In: Croft, D., Su, D., Simpson, S. (eds) Methods in Paleoecology. Vertebrate Paleobiology and Paleoanthropology. Springer, Cham. doi: 10.1007/978-3-319-94265-0_2

Huaiyu, M., Muruganujan, A., Huang, J. X., Ebert, D., Mills, C., Guo, X. & Thomas, P. D. (2019). Protocol Update for large-scale genome and gene function analysis with the PANTHER classification system (v.14.0). Nature Protocols 14, 703–721. doi: 10.1038/s41596-019-0128-8

Ishida, K., Murofushi, M., Nakao, K., Morita, R., Ogawa, M., & Tsuji, T. (2011). The regulation of tooth morphogenesis is associated with epithelial cell proliferation and the expression of Sonic hedgehog through epithelial-mesenchymal interactions. Biochemical and Biophysical Research Communications, 405(3), 455–461. doi: 10.1016/j.bbrc.2011.01.052

Jia, S., Zhou, J., Gao, Y., Baek, J.-A., Martin, J. F., Lan, Y. & Jiang, R. (2013). Roles of Bmp4 during tooth morphogenesis and sequential tooth formation. Development, 140, 423–432. doi: 10.1242/dev.081927

Jing, J., Feng, J., Yan, Y., Guo, T., Lei, J., Pei, F., Ho, T.-V. & Chai, Y. (2022). Spatiotemporal single-cell regulatory atlas reveals neural crest lineage diversification and cellular function during tooth morphogenesis. Nature Communications, 13, 4803. doi: 10.1038/s41467-022-32490-y

Jernvall, J. (1995). Mammalian molar cusp patterns: Developmental mechanisms of diversity. Acta Zoologica Fennica, 198. 1–61.

Jernvall, J., Keränen, S. V. E., & Thesleff, I. (2000). Evolutionary modification of development in mammalian teeth: Quantifying gene expression patterns and topography. Proceedings of the National Academy of Sciences, 97(26), 14444–14448. doi: 10.1073/pnas.97.26.14444

Joseph, B. K., Savage, N. W., Daley, T. J., & Young, W. G. (1996). In Situ Hybridization Evidence for a Paracrine/Autocrine Role for Insulin-Like Growth Factor-I in Tooth Development. Growth Factors, 13(1-2), 11–17. doi: 10.3109/08977199609034563

Järvinen, E., Salazar-Ciudad, I., Birchmeier, W., Taketo, M. M., Jernvall, J., & Thesleff, I. (2006). Continuous tooth generation in mouse is induced by activated epithelial Wntbeta-catenin signaling. Proceedings of the National Academy of Sciences, 103(49), 18627–18632. doi: 10.1073/pnas.0607289103

Järvinen, E., Välimäki, K., Pummila, M., Thesleff, I., & Jernvall, J. (2008). The taming of the shrew milk teeth. Evolution & Development, 10(4), 477–486. doi: 10.1111/j.1525-142x.2008.00258.x.

Kinsella, R. J., Kähäri, A., Haider, S., Zamora, J., Proctor, G., Spudich, G., … Flicek, P. (2011). Ensembl BioMarts: a hub for data retrieval across taxonomic space. Database, 2011(0), bar030. doi: 10.1093/database/bar030

Koffi, K. A., Doublier, S., Ricort, J.-M., Babajko, S., Nassif, A. & Isaac, J. (2021). The Role of GH/IGF Axis in Dento-Alveolar Complex from Development to Aging and Therapeutics: A Narrative Review. Cells, 10, 1181. doi.org/10.3390/cells10051181

Kopylova, E., Noé, L., & Touzet, H. (2012). SortMeRNA: fast and accurate filtering of ribosomal RNAs in metatranscriptomic data. Bioinformatics, 28(24), 3211–3217. doi: 10.1093/bioinformatics/bts611

Kuiper, S. & Sklar, J. (2012). Practicing statistics: Guided investigations for the second course. Pearson Higher Ed.

Lafuma, F., Corfe, I. J., Clavel, J., & Di-Poï, N. (2021). Multiple evolutionary origins and losses of tooth complexity in squamates. Nature Communications, 12(1), 6001. doi: 10.1038/s41467-021-26285-w

LeRoith, D., Holly J. M. P. & Forbes, B. E. (2021). Insulin-like growth factors: Ligands, binding proteins, and receptors. Molecular Metabolism, 52, 101245. doi: 10.1016/j.molmet.2021.101245

Larionova, D., Lesot, H. & Huysseune, A. (2021). Miniaturization: How many cells are needed to build a tooth? Developmental Dynamics 250, 1021–1035. doi: 10.1002/dvdy.300

Love, M. I., Huber, W., & Anders, S. (2014). Moderated estimation of fold change and dispersion for RNA-seq data with DESeq2. Genome Biology, 15(12), 550. doi: 10.1186/s13059-014-0550-8

Luo, Z.-X. (2007). Transformation and diversification in early mammal evolution. Nature, 450(7172), 1011–1019. doi: 10.1038/nature06277

Metscher, B. D. (2009). MicroCT for comparative morphology: simple staining methods allow high-contrast 3D imaging of diverse non-mineralized animal tissues. BMC Physiol 9, 11. doi.org/10.1186/1472-6793-9-11

Mogollón, I., Moustakas-Verho, J. E., Niittykoski, M., & Ahtiainen, L. (2021). The initiation knot is a signaling center required for molar tooth development. Development, (9). doi: 10.1242/dev.194597

Nasrullah, G., Renfree, M. & Evans, A. R. (2022). From embryo to adult: the complete development and unusual replacement of the dentition of the tammar wallaby (Macropus eugenii). Journal of Mammalian Evolution, 29, 515-529. doi.org/10.1007/s10914-021-09597-y

Närhi, K., & Thesleff, I. (2010). Oral Biology, Molecular Techniques and Applications. Methods in Molecular Biology, 666, 253–267. doi: 10.1007/978-1-60761-820-1_16

Ohyama, T. & Groves, A. K. (2004). Expression of mouse Foxi class genes in early craniofacial development. Developmental Dynamics 231(3), 640–646. doi: 10.1002/dvdy.20160.

Oyanagi, T., Takeshita, N., Hara, M., Ikeda, E., Chida, T., Seki, D., … Takano-Yamamoto, T. (2019). Insulin-like growth factor 1 modulates bioengineered tooth morphogenesis. Scientific Reports, 9(1), 368. doi: 10.1038/s41598-018-36863-6

Pampush, J. D., Winchester, J. M., Morse, E., Paul, A. Q., Vining, D. M., Boyer, Doug & R. F., Kay (2016). Introducing molaR: a new R package for quantitative topographic analysis of teeth (and other topographic surfaces). Journal of Mammalian Evolution 23(4), 397–412. doi: 10.1007/s10914-016-9326-0

Parker, J. (2011). Morphogens, nutrients, and the basis of organ scaling. Evolution & Development, 13(3), 304–314. doi: 10.1111/j.1525-142x.2011.00481.x

Patro, R., Duggal, G., Love, M. I., Irizarry, R. A., & Kingsford, C. (2017). Salmon provides fast and bias-aware quantification of transcript expression. Nature Methods, 14(4), 417–419. doi: 10.1038/nmeth.4197

Peters, R. (1983). The Ecological Implications of Body Size. Cambridge: Cambridge University Press. doi:10.1017/CBO9780511608551

Rainer, J., Gatto, L., & Weichenberger, C. X. (2019). ensembldb: an R package to create and use Ensembl-based annotation resources. Bioinformatics, 35(17), btz031. doi: 10.1093/bioinformatics/btz031

Renvoisé, E., Kavanagh, K. D., Lazzari, V., Häkkinen, T. J., Rice, R., Pantalacci, S., … Jernvall, J. (2017). Mechanical constraint from growing jaw facilitates mammalian dental diversity. Proceedings of the National Academy of Sciences, 114(35), 9403–9408. doi: 10.1073/pnas.1707410114

Sakaue-Sawano, A., Kurokawa, H., Morimura, T., Hanyu, A., Hama, H., Osawa, H., … Miyawaki, A. (2008). Visualizing Spatiotemporal Dynamics of Multicellular Cell-Cycle Progression. Cell, 132(3), 487–498. doi: 10.1016/j.cell.2007.12.033

Salazar-Ciudad, I. & Jernvall, J. (2010). A computational model of teeth and the developmental origins of morphological variation. Nature 464, 583–586. doi: 10.1038/nature08838

Sasaki, T., Ito, Y., Xu, X., Han, J., Bringas Jr. P., Maeda, T., Slavkin, H. C., Grosschedl, R. & Chai, Y. (2005). LEF1 is a critical epithelial survival factor during tooth morphogenesis. Developmental Biology, 278, 130–143. doi.org/10.1016/j.ydbio.2004.10.021

Savriama, Y., Valtonen, M., Kammonen, J. I., Rastas, P., Smolander, O.-P., Lyyski, A., … Jernvall, J. (2018). Bracketing phenogenotypic limits of mammalian hybridization. Royal Society Open Science, 5(11), 180903. doi: 10.1098/rsos.180903

Schindelin, J., Arganda-Carreras, I., Frise, E., Kaynig, V., Longair, M., Pietzsch, T., … Cardona, A. (2012). Fiji: an open-source platform for biological-image analysis. Nature Methods, 9(7), 676–682. doi: 10.1038/nmeth.2019

Shirokova, V., Jussila, M., Hytönen, M. K., Perälä, N., Drögemüller, C., Leeb, T., … Mikkola, M. L. (2013). Expression of Foxi3 is regulated by ectodysplasin in skin appendage placodes. Developmental Dynamics, 242(6), 593–603. doi: 10.1002/dvdy.23952

Sjögren, K., Liu, J.-L., Blad, K., Skrtic, S., Vidal, O., Wallenius, V., … Ohlsson, C. (1999). Liver-derived insulin-like growth factor I (IGF-I) is the principal source of IGF-I in blood but is not required for postnatal body growth in mice. Proceedings of the National Academy of Sciences, 96(12), 7088–7092. doi: 10.1073/pnas.96.12.7088

Smith, F. A., Boyer, A. G., Brown, J. H., Costa, D. P., Dayan, T., Ernest, 4 S. K. Morgan, … Uhen, M. D. (2010). The Evolution of Maximum Body Size of Terrestrial Mammals. Science, 330(6008), 1216–1219. doi: 10.1126/science.1194830

Smith, F. A., Payne, J. L., Heim, N. A., Balk, M. A., Finnegan, S., Kowalewski, M., … Wang, S. C. (2016). Body Size Evolution Across the Geozoic. Annual Review of Earth and Planetary Sciences, 44(1), 1–31. doi: 10.1146/annurev-earth-060115-012147

Sokal, R. R. & Rohlf, F. J. (1995). Biometry (Freeman, New York).

Soneson, C., Love, M. I., & Robinson, M. D. (2016). Differential analyses for RNA-seq: transcript-level estimates improve gene-level inferences. F1000Research, 4, 1521. doi: 10.12688/f1000research.7563.2

Sonnhammer, E. L. L., & Östlund, G. (2015). InParanoid 8: orthology analysis between 273 proteomes, mostly eukaryotic. Nucleic Acids Research, 43(D1), D234–D239. doi: 10.1093/nar/gku1203

Streelman, J. T., Webb, J. F., Albertson, R. C., & Kocher, T. D. (2003). The cusp of evolution and development a model of cichlid tooth shape.pdf. Evolution and Development, 5(6), 600–608. doi: https://doi.org/10.1046/j.1525-142X.2003.03065.x

Sutter, N. B., Bustamante, C. D., Chase, K., Gray, M. M., Zhao, K., Zhu, L., … Ostrander, E. A. (2007). A Single IGF1 Allele Is a Major Determinant of Small Size in Dogs. Science, 316(5821), 112–115. doi: 10.1126/science.1137045

Thiery, A. P., Standing, A. S., Cooper, R. L., & Fraser, G. J. (2022). An epithelial signalling centre in sharks supports homology of tooth morphogenesis in vertebrates. ELife, 11, e73173. doi: 10.7554/elife.73173

Thomas, P. D., Ebert, D., Muruganujan, A., Mushayahama, T., Albou, L.-P. & Huaiyu, M. (2022). PANTHER: Making genome-scale phylogenetics accessible to all. Protein Society 31, 8–22. doi:10.1002/pro.4218

Wilkinson, D. G., & Nieto, M. A. (1993). Detection of Messenger RNA by in Situ Hybridization to Tissue Sections and Whole Mounts. Methods in Enzymology, 225, 361–373. doi: 10.1016/0076-6879(93)25025-w.

Winchester, J. M. (2016). MorphoTester: An Open Source Application for Morphological Topographic Analysis. PLoS ONE 11(2): e0147649. doi.org/10.1371/journal.pone.0147649

Woods, K. A., Camacho-Hübner, C., Savage, M. O., & Clark, A. J. L. (1996). Intrauterine growth retardation and postnatal growth failure associated with deletion of the insulin-like growth factor I gene. The New England Journal of Medicine, 355(18), 1363–1367. doi: 10.1056/NEJM199610313351805.

Wu, Z., & Guan, K.-L. (2020). Hippo Signaling in Embryogenesis and Development. Trends in Biochemical Sciences, 46(1), 51–63. doi: 10.1016/j.tibs.2020.08.008

Young, W.G., Ruch, J.V., Stevens, M.R., Bègue-Kirn, C., Zhang, C.Z., Lesot, H. & WatersM J. (1995) Comparison of the effects of growth hormone, insulin-like growth factor-I and fetal calf serum on mouse molar odontogenesis in vitro. Archives of Oral Biology, 40(9):789–99. doi: 10.1016/0003-9969(95)00051-p.

